# Optogenetic Stimulation of Pancreatic Function via Vagal Cholinergic Axons

**DOI:** 10.1101/2019.12.21.885970

**Authors:** Arjun K. Fontaine, David G. Ramirez, Samuel F. Littich, Robert A. Piscopio, Vira Kravets, John H. Caldwell, Richard F. Weir, Richard K.P. Benninger

## Abstract

Previous studies have demonstrated stimulation of endocrine pancreas function by vagal nerve electrical stimulation. While this increases insulin secretion; concomitant reductions in circulating glucose do not occur. A complicating factor is the non-specific nature of electrical nerve stimulation. Optogenetic tools enable high specificity in neural stimulation using cell-type specific targeting of opsins and/or spatially shaped excitation light. Here, we demonstrate light-activated stimulation of the endocrine pancreas by targeting vagal parasympathetic axons. In a mouse model expressing ChannelRhodopsin2 (ChR2) in cholinergic cells, serum insulin and glucose were measured in response to both ultrasound image-guided optical stimulation of axon terminals in the pancreas and optical stimulation of axons of the cervical vagus nerve, together with ultrasound-based measures of pancreas blood flow. Measurements were made in basal-glucose and glucose-stimulated conditions. Significant increases in plasma insulin occurred relative to controls under both pancreas and vagal stimulation, accompanying rapid reductions in glycemic levels. Additionally, a significant increase in pancreatic blood flow was measured following optical stimulation. Together, these results demonstrate the utility of in-vivo optogenetics for studying the neural regulation of endocrine pancreas function and suggest therapeutic potential for the control of insulin secretion and glucose homeostasis.

---

Neural modulation of the endocrine pancreas is important for insulin secretion and glucose homeostasis. Vagus nerve electrical stimulation stimulates insulin secretion in multiple mammalian animal models^1–6^. Parasympathetic (cholinergic) inputs mediate this effect, as it is reduced or abolished with acetylcholine receptor antagonists^1,2,4,5^. While differences exist, cholinergic inputs are observed within both mouse islet and human islet^7,8^.

Enhancing insulin secretion is a central goal in treating patients with Type 2 Diabetes^9^. Side effects with existing therapies (hypoglycemia and beta cell apoptosis with K_ATP_ inhibitors^9,10^; nausea and pancreatitis with GLP1R agonists^11^) suggest other targets should be considered. A temporally controlled and precisely targeted approach to increase insulin secretion would be advantageous.

Neural interfacing, particularly at the peripheral nerve level, is increasingly studied and utilized as a mode of organ modulation for disease treatment^12,13^. Vagus nerve stimulation therapies have shown efficacy across numerous clinical areas, including inflammation in rheumatoid arthritis^14^, epileptic seizures^15^, depression^16^, obesity^17^, and migraine^18^. In many cases, however, the underlying neural mechanisms are poorly understood, in part because the electrical stimulation lacks axon-level specificity. Thus, activation of off-target pathways can occur. For example, in animal models, while insulin is increased with electrical vagal stimulation, glucose remains unchanged or even increases^1–3,6^. Given the established understanding that insulin acts to decrease circulating glucose, this demonstrates that counteracting (off-target) pathways, such as glucagon activation or its targets, may interfere with a signaling pathway of interest.

Highly specific activation of neural pathways is possible by means of optogenetics, whereby a light-activatable ion channel or pump can be genetically targeted to a specific cell type. While this optical approach is genetically defined, it can achieve further specificity based on spatial shaping of the excitation light^19^. Here, we demonstrate light-activated stimulation of the endocrine pancreas by targeting vagal parasympathetic axons. Using a mouse model expressing ChannelRhodopsin2 (ChR2) in cholinergic cells we demonstrate increases in plasma insulin, rapid decreases in blood glucose and increased blood flow in response to optical stimulation of axon terminals at the pancreas and axons in the cervical vagus nerve.

## Research Design and Methods

### Animals

All animal procedures were performed in accordance with guidelines established by the Institutional Animal Care and Use Committee of the University of Colorado. Double homozygous ChAT-Cre; Rosa-ChR2 mice (‘ChAT-ChR2’) were bred in house. Age-matched wild-type C57Bl6 mice (JAX) were used as controls. Mice were fasted 6h prior to each experiment.

### Direct Pancreas Stimulation

Mice were anesthetized using 1-3% isoflurane, and placed supine on a heating pad and monitored throughout the experiment for body temperature and vitals measurements. Cannula placement was guided by ultrasound imaging with a VEVO2100 small animal ultrasound machine (Visual Sonics). The pancreas was identified by location in relation to the spleen, kidney, and stomach, and by image texture. An optical cannula (CFMC12L20, Thorlabs) was guided into the abdomen within a 22G needle and directed to the pancreas. A 473nm-wavelength solid-state laser (SLOC) was coupled to the optical cannula with an optical patch cable (200µm core, .39 NA, M81L005, Thorlabs) and an interconnect (ADAF2, Thorlabs). The laser was controlled with a custom Arduino circuit board to output 5ms pulses at 20 Hz. The continuous-wave (non-pulsed) power output at the tip of the cannula was 45 mW. Blood glucose was measured from tail vein samples using a glucose meter (Bayer Contour) every five minutes throughout the experiment. After a 20min baseline period, laser stimulation of the pancreas was applied for 25 minutes. Immediately following the laser stimulation blood was collected into a heparin-coated collection tube, and insulin concentration assessed by ELISA (Alpco). For glucose tolerance testing, a 200mg/ml glucose solution in PBS was injected intraperitoneally at a volume of 10µl/g bw. Following stimulation, mice were monitored until full recovery.

### Blood Flow Measurements

Blood flow in the pancreas was measured using contrast-enhanced ultrasound (CEUS) imaging, as previously described^20^. Briefly, acquisition settings of the MS250 linear array transducer were set at 10% transmit power, 18MHz, standard beamwidth, contrast gain of 30dB, 2D gain of 18dB, with an acquisition rate of 26fps. Microbubbles (3-4µm size-isolated, Advanced Microbubbles Laboratories) were injected as a single 100µl bolus (10×10^6^ MBs), into the lateral tail via a custom-made 27G½” winged infusion set. Following infusion, contrast intensity was measured and allowed to reach a steady state. A high mechanical-index flash-destruction was then initiated within a region of interest of the pancreas (identified using B-mode imaging, see above). Recovery of contrast signal within this region was measured as a time-course averaged over the pancreas, normalized to the contrast signal over the last five seconds prior to flash-destruction. The rise rate of the reperfusion was determined through an exponential fit.

### Cervical Vagus Stimulation

Vitals were monitored with a MouseOx Plus suite (Starr Life Sciences). A 1-1.5cm incision was made at the cervical region, 2-3mm left of midline. Blunt dissection techniques were used to expose the left cervical vagus nerve and separate it from the carotid artery. The optical cannula was positioned with a micromanipulator to abut the nerve for laser stimulation. Baseline glucose measurements were obtained as described above, once the surgical opening was complete. The laser pulsing was the same as in the direct pancreas stimulation (5ms pulse duration, at 20Hz).

The continuous-wave (non-pulsed) power at the output of the cannula was 36mW. A small downward slope in glucose during the procedure was slope-detrended using the linear regression of the non-laser-stimulated ChAT-ChR2 group. This slope was subtracted from all three experimental groups in Figure 3.

**Figure 1:**
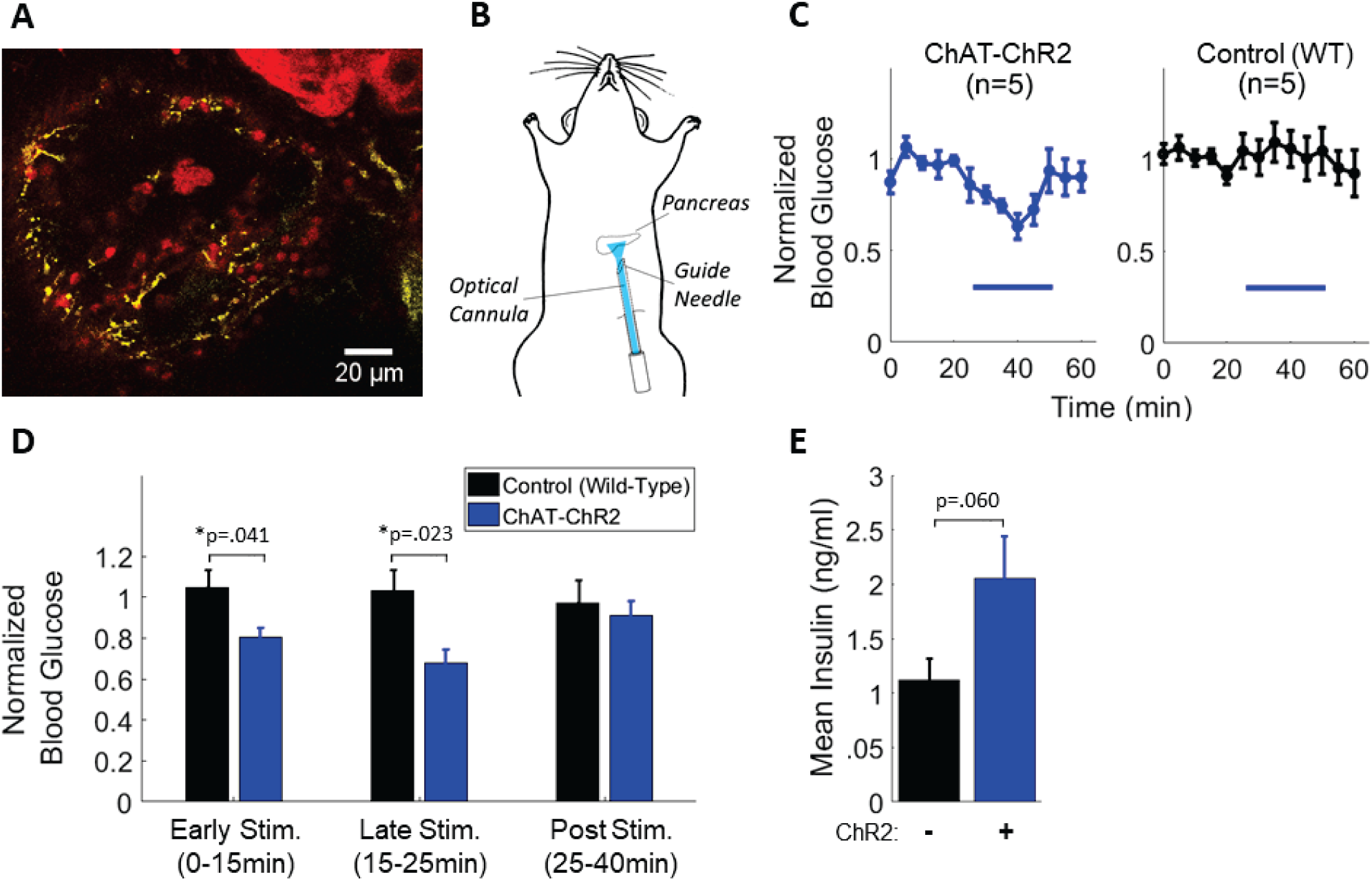
Direct optogenetic control of pancreas innervation at basal glucose. (A) Maximum projection image of a pancreatic islet within a tissue slice from a ChAT-ChR2 mouse, showing YFP fluroescence indicating regions of ChR2 expression in axon/terminals (yellow), and surrounding tissue labeled with Rhod-2 AM (red). (B) Diagram of optical stimulation at the pancreas in the anesthetized mouse. (C) Blood glucose in ChAT-ChR2 mice and wild-type mice experimental groups in response to optical stimulation (blue bar, 473nm, 5 ms pulses, 20 Hz), and following stimualtion. (D) Quantification of blood glucose levels during the early stimulation period (0-15min from onset of stimulation), late stimulation period (15-25min), and post stimulation period (25-40min) in ChAT-ChR2 mice and wild-type mice. * Indicates statistically significant difference by student’s t-test. (E) Post-stimulation mean insulin levels of the two experimental groups sampled immediately following laser stimulation.

**Figure 2:**
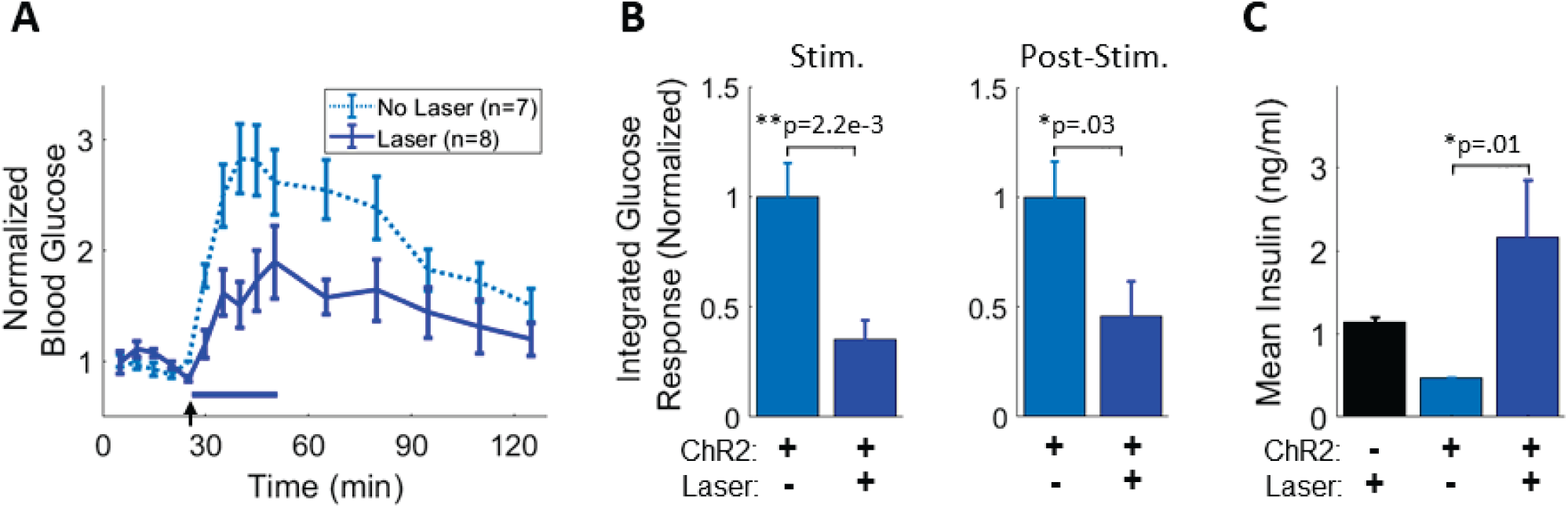
Direct optogenetic control of pancreas innervation following glucose elevation. (A) Blood glucose response in ChAT-ChR2 mice with and without laser stimulation (blue bar, 473nm, 5 ms pulses, 20 Hz) following intraperitoneal glucose bolus injection (black arrow). (B) Integrated area under the glucose response curve during the stimulation period (left) and during the post-stimulation period (right). (C) Blood insulin concentration sampled immediately following stimulation is significantly increased in the laser-stimulated ChAT-ChR2 group relative to non-laser-stimulated ChAT-ChR2.

**Figure 3:**
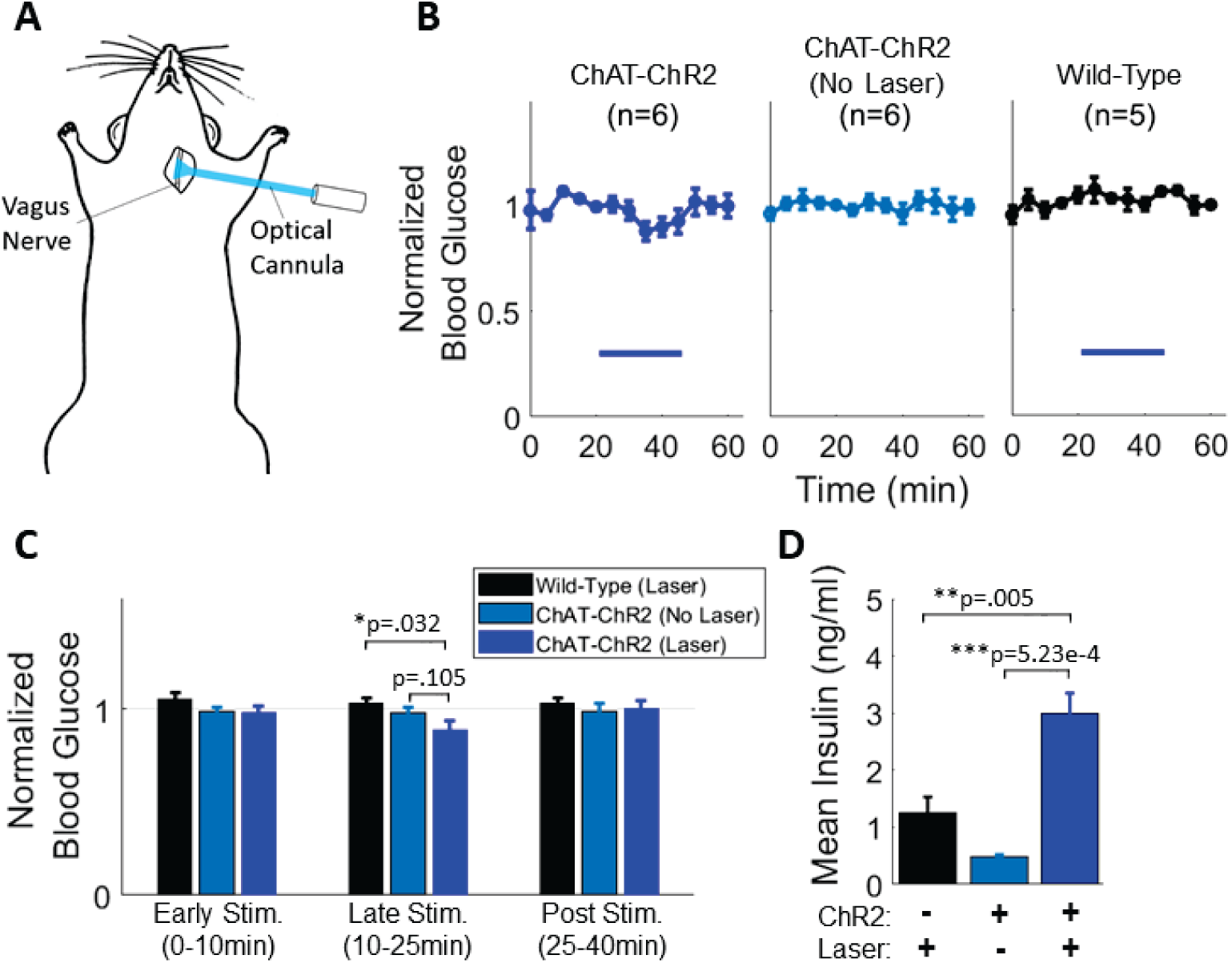
Cervical vagus nerve stimulation in the basal glucose state. (A) Schematic diagram of the location of laser stimulation. (B) Blood glucose levels in laser-stimulated ChAT-ChR2 mice (left), non-laser-stimulated ChAT-ChR2 mice (middle) and laser-stimulated wild-type mice (right)(laser stimulation indicated by blue bar, 5 ms pulses, 20 Hz). A decrease in glucose is observed in the optically stimulated ChAT-ChR2 group. (C) Quantification of glucose levels in the early stimulation period (0-10min following stimulation onset), late stimulation period (10-25min) and post stimulation period (25-40min). (D) Quantification of post stimulation insulin levels showed a large increase in the laser-stimulated ChAT-ChR2 group relative to controls.

## Results

### Direct pancreas cholinergic stimulation impacts basal glucose homeostasis

We first tested whether insulin and glucose could be modulated by direct pancreatic stimulation of cholinergic axons. Confocal imaging of YFP in pancreas slices from ChAT-ChR2 mice confirmed the presence of ChR2-expressing cholinergic axons that terminated within islets (Figure 1A). In fasted mice (6h), an optical cannula was ultrasound-guided to the pancreas (Figure 1B). Glucose levels in ChAT-ChR2 mice rapidly decreased upon laser stimulation and recovered to baseline following stimulation. Wild-type control mice did not exhibit stimulation-dependent decreases (Figure 1C). Blood glucose in ChAT-ChR2 mice was significantly lower than in wild-type controls, both early (0-15min, -23%, p=.04) and late (15-25min, -34%, p=.02) in the stimulation period but returned to control levels after laser stimulation (Figure 1D). Insulin levels in the ChAT-ChR2 group were higher (+84%, p=.06) than in the wild-type controls after laser stimulation (Figure 1E).

### Direct pancreas cholinergic stimulation affects glucose excursions

We next applied optogenetic stimulation to the pancreas during acute glucose elevations. Following glucose delivery, glycemic levels were significantly blunted during (−65%, p=.002) and after (−54%, p=.03) laser stimulation relative to non-laser-stimulated ChAT-ChR2 mice (Figure 2A,B). Insulin levels were also significantly increased in laser-stimulated ChAT-ChR2 mice compared to non-laser-stimulated ChAT-ChR2 mice (+363%, p=.01)(Figure 2C). ChAT-ChR2 mice show elevated glucose excursions relative to wild-type mice (Supplementary Figure 1) preventing a comparison between laser-stimulated ChAT-ChR2 and control mice.

### Cervical Vagus cholinergic stimulation impacts glucose homeostasis

To test whether similar modulation of insulin secretion and glucose homeostasis could be achieved by proximal nerve stimulation, we stimulated cholinergic axons within the cervical vagus nerve. Mice were surgically opened in the cervical region to allow vagus nerve exposure and optical stimulation (Figure 3A). Laser stimulation of the left cervical vagus nerve reduced blood glucose in ChAT-ChR2 mice but not in control mice (Figure 3B). Blood glucose in ChAT-ChR2 mice was significantly lower than in wild-type controls late (15-25min, -14%, p=.032) in the stimulation period and returned to control levels after laser stimulation (Figure 3C). Insulin levels in ChAT-ChR2 were higher than in the wild-type controls (+140%, p=.005) and non-laser stimulated ChAT-ChR2 mice (+512%, p=.0005) (Figure 3D).

### Cholinergic stimulation increases pancreatic blood flow

Parasympathetic activity has been suggested to modulate pancreas and islet blood flow^21,22^, therefore we next tested whether cholinergic stimulation would modulate pancreas blood flow. We quantified pancreas vascular perfusion dynamics using contrast-enhanced ultrasound following direct pancreas laser stimulation in 6h fasted mice (Figure 4A). Following flash-destruction of microbubbles within the pancreas, their recovery kinetics were measured before and after laser stimulation (Figure 4B). The mean reperfusion rate increased significantly in ChAT-ChR2 mice (+123%, p=.05) indicating more rapid blood flow, while the wild-type group had no change in reperfusion rate following stimulation (p=.56) (Figure 4C).

**Figure 4:**
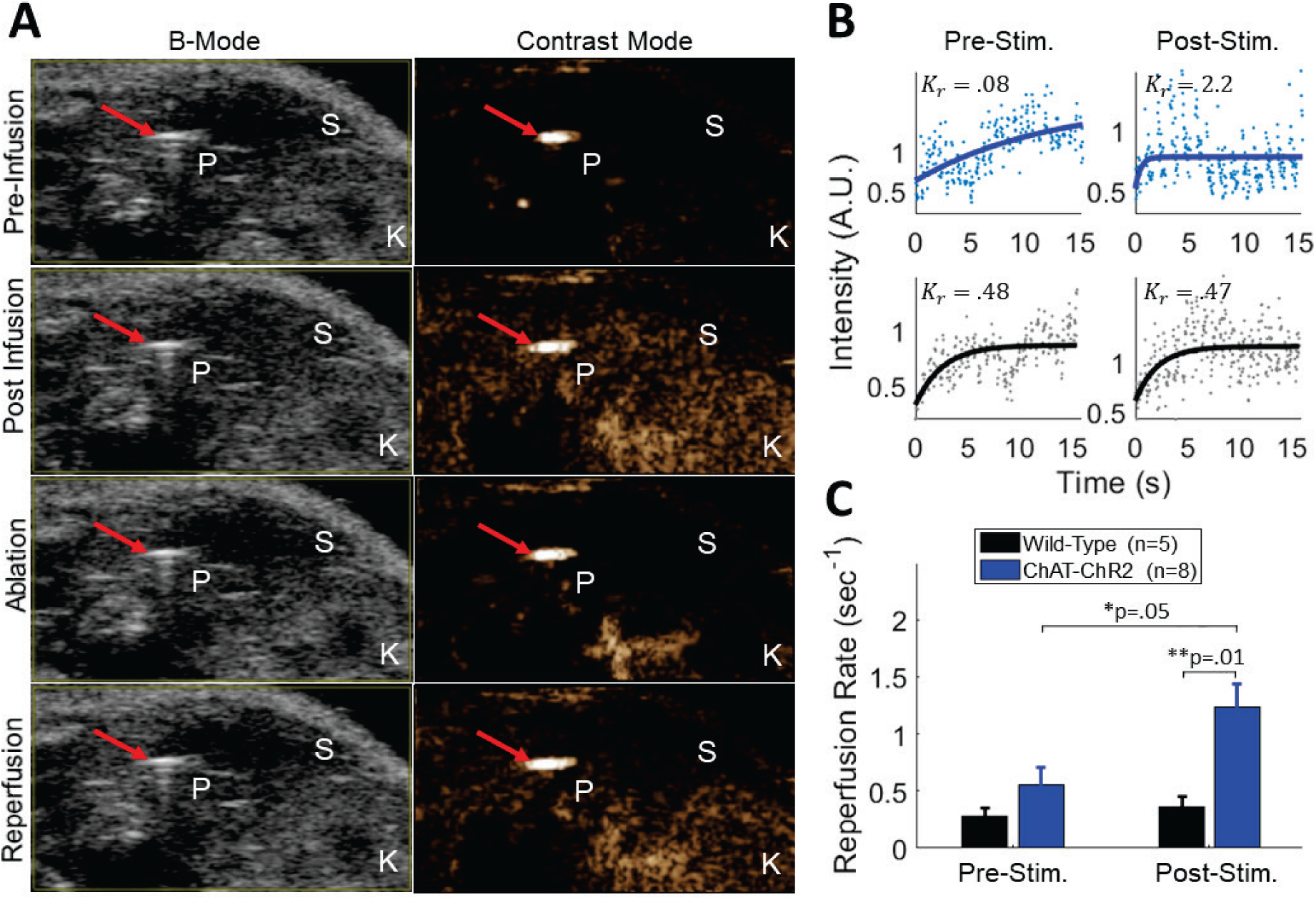
Measurement pancreatic blood perfusion using contrast enhanced ultrasound imaging of microbubbles. (A) B-Mode and sub-harmonic contrast mode images in the abdominal cavity. Microbubble-generated sub-harmonic contrast is observed in the tissues during microbubble infusion and during reperfusion following flash-destruction. Red arrow = optical cannula/needle tip, P = pancreas (tail), S = spleen, K = kidney. (B) Example single recording reperfusion data and curve fit before and after laser stimulation for a ChAT-ChR2 mouse and wild-type mouse. Intensity is based on microbubble (MB)-generated sub-harmonic contrast as MBs perfuse back into the pancreas following flash destruction. Curves are fitted to a single component inverse exponential to extract reperfusion rate (K_r_). Upper panels are from a ChAT-ChR2 mouse and lower panels are from a wild-type mouse. (C) Mean reperfusion rate in ChAT-ChR2 mice and wild-type mice, prior to laser stimulation and after 25minutes of laser stimulation.

## Discussion

Electrical stimulation of the vagus nerve can stimulate endocrine pancreas hormone secretion; particularly insulin secretion. However reductions in blood glucose are not generally achieved ^1–3,5,6^. The relatively indiscriminant electrical stimulation likely activates off-target pathways such as sensory fibers and fibers that innervate other organs. In the present study, we employed an optogenetic technique which enabled the optical stimulation of only parasympathetic (and thus primarily efferent) vagal fibers via expression of ChR2 in cholinergic axons. With this optical approach, insulin secretion was elevated in response to stimulation at the pancreas and cervical vagus nerve. Importantly, and in contrast to electrical stimulation approaches, blood glucose was rapidly reduced during the optical stimulation, under both basal glucose and glucose-elevated conditions. Blood flow was also increased in the pancreatic vasculature.

As vagal cholinergic axons are broadly efferent, the likely explanation for the robust glucose reduction under optical stimulation is the omission of afferent fiber activation that would occur under electrical stimulation. Vagus nerve afferent stimulation does increase blood glucose in rats^6^. Furthermore, hepatic vagal afferents have been suggested to inhibit the brainstem centers which drive the efferent system that controls insulin secretion^23^. Thus, the afferent activation during electrical stimulation of the whole vagus nerve may dampen insulin action, resulting in higher blood glucose levels.

Interestingly, we observed a more pronounced reduction in blood glucose during direct pancreas stimulation compared to cervical vagus stimulation. This occurred despite insulin increasing more robustly under cervical stimulation, and thus not as a result of less efficient nerve stimulation. The left vagus nerve innervates almost all the thoracic and abdominal organs, and thus cervical stimulation may be activating other organs that impact glucose homeostasis which are not activated under direct pancreas stimulation. This may include reducing insulin action and glucose uptake, or stimulating hepatic glucose production. Alternatively, stimulation at the vagus may have preferentially stimulated more glucagon secretion, which can be activated by cholinergic inputs to the pancreas^1,4^. Nevertheless, these results also suggest that electrical stimulation may be directly activating other efferent pathways that reduce insulin action and prevent efficient modulation of glycemia. This points to the advantage provided by optogenetic approaches and image-guided stimulation to study and control neural inputs at the organ level.

The observed increase in blood flow within the pancreas are expected from parasympathetic stimulation. Cholinergic stimulation has been suggested (but not quantitatively demonstrated) to increase blood flow in the pancreas. Blocking cholinergic receptors with atropine caused ischemia in pancreatic capillary beds^21^. Under increased glucose levels, during which parasympathetic activity increases, more rapid islet blood flow occurs relative to decreased glucose levels^22,24^. Conversely, stimulation of adrenergic nerve fibers decreases pancreatic blood flow^5,24^. The increase in blood flow under parasympathetic activity may contribute to increased insulin release, although precise links between islet blood flow and hormone secretion remain to be determined.

The results of this study, utilizing novel optogenetic and image-guiding approaches, demonstrate effective control of both hormone secretion and blood glucose levels, which is not observed with whole nerve electrical stimulation. This highlights the functional improvement that can be achieved by increasing neural specificity, and the potential of optogenetic interfaces towards this. This methodology will be valuable for studying signaling pathways within the endocrine pancreas *in vivo*, particularly those involving cellular excitability. While the reduction in blood glucose in response to cervical vagus stimulation was modest, despite robust insulin secretion, the efficacy of cervical stimulation may be improved by furthering the specificity of opsin expression to pancreas-specific axons. Future studies should therefore investigate the use of a pancreas-injected retrograde adeno-associated virus (AAV)^25^ to transduce opsin expression in appropriate pancreatic axons. Additional steps toward a potential clinically translatable therapy include the integration of an implanted nerve cuff to allow stimulation in awake and behaving subjects. Combined with continuous glucose monitoring and closed-loop control, effective glycemic control may be achieved.

## Acknowledgements

This work was supported by the NIH Office of the Director, SPARC Initiative, grant 3OT2OD023852-01S4 (Caldwell, Gibson, Weir): ‘Development of a Bidirectional Optogenetic Minimally Invasive Peripheral Nerve Interface with Single Axon Read-in & Read-out Specificity’, as well as NIH grants R01 DK102950, R01 DK106412; and Juvenile Diabetes Research Foundation Grant 5-CDA-2014-198-A-N (to RKPB). Engineering support was provided by the Optogenetics and Neural Engineering Core at the University of Colorado Anschutz Medical Campus, funded in part by the National Institute of Neurological Disorders and Stroke of the NIH under award number P30 NS048154.

A.K.F. designed and performed experiments, analyzed the data, and wrote the manuscript. D.G.R. and A.K.F. performed the direct pancreas stimulation experiments, with D.G.R. conducting the ultrasound imaging and microbubble-based blood flow measurements. A.K.F. performed the cervical vagus nerve stimulation surgical procedures. S.L. assisted the cervical vagus experiments. R.P. performed Elisa assays and V.K. and R.P. prepared pancreas slices for imaging. J.H.C. and R.F.W. provided advising and research support and edited the manuscript. R.K.P.B. provided advising and research support, designed experiments, and edited the manuscript.

The authors declare no conflicts of interest.

## Supplemental Figures

**Supplementary Figure 5:**
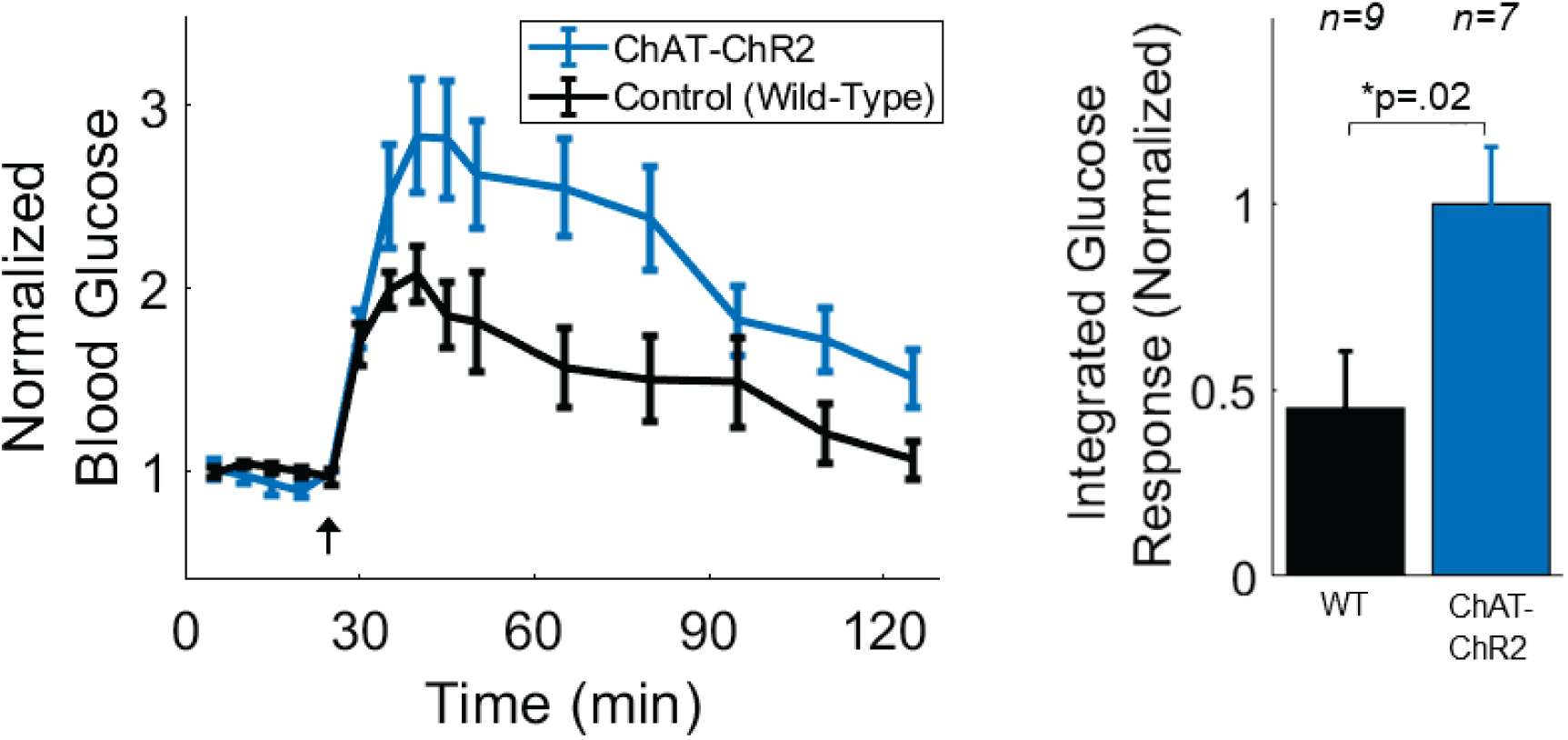
Glucose tolerance tests in ChAT-ChR2 and wild-type mice with no laser stimulation. (Left) Glucose measurements following bolus glucose injection (black arrow) show increased glucose response amplitude in ChAT-ChR2 mice relative to wild-type control mice. (Right) Integrated response under the glucose response curve is significantly larger in ChAT-ChR2 mice compared to wild-type mice.

